# Yeast Ssd1 is a non-enzymatic member of the RNase II family with an alternative RNA recognition interface

**DOI:** 10.1101/2020.10.22.350314

**Authors:** Rosemary A. Bayne, Uma Jayachandran, Aleksandra Kasprowicz, Stefan Bresson, David Tollervey, Edward W. J. Wallace, Atlanta G. Cook

## Abstract

The conserved fungal RNA binding protein Ssd1, is important in stress responses, cell division and virulence. Ssd1 is closely related to Dis3L2 of the RNase II family of nucleases, but lacks catalytic activity and may act by suppressing translation of associated mRNAs. Previous studies identified motifs that are enriched in Ssd1-associated transcripts, yet the sequence requirements for Ssd1 binding are not well understood. Here we present the crystal structure of Ssd1 at 1.9 Å resolution. Active RNase II enzymes have a characteristic, internal RNA binding path, but in Ssd1 this is blocked by remnants of regulatory sequences. Instead, RNA binding activity has likely been relocated to the outer surface of the protein. Using *in vivo* crosslinking and cDNA analysis (CRAC), we identify Ssd1-RNA binding sites. These are strongly enriched in 5’UTRs of a subset of mRNAs encoding cell wall proteins. Based on these and previous analyses, we identified a conserved bipartite motif that binds Ssd1 with high affinity *in vitro*. These studies provide a new framework for understanding the function of a pleiotropic post-transcriptional regulator of gene expression and give insights into the evolution of regulatory elements in the RNase II family.

## Introduction

Mechanisms of post-transcriptional control of gene expression by RNA binding proteins (RBPs) include modulation of mRNA translation and decay. The RNase II/RNB family enzymes are found in all domains of life, where they play roles in RNA maturation and degradation (Reis et al. 2013). Eukaryotic DIS3 (Rrp44) and Dis3L2 are RNB family 3’-5’ exonucleases. DIS3 or Rrp44 (for human and yeast orthologues respectively) is the essential nuclease associated with the eukaryotic exosome complex that processes and/or turns over the majority of cellular RNAs (Dziembowski et al. 2007). Dis3L2 is a related nuclease that is specific for RNA substrates with an oligouridine 3’ tail (Malecki et al. 2013). However, some RNase II family proteins are pseudonucleases with regulatory roles in RNA metabolism, rather than active enzymes. These include the fungal Ssd1 family that is closely related to Dis3L2 (Ballou, Cook, and Wallace 2020). Ssd1 was initially identified in *Saccharomyces cerevisiae* as a genetic suppressor of mutations in the Sit4 protein phosphatase (Sutton, Immanuel, and Arndt 1991). While no exonuclease activity could be detected from Ssd1, RNA binding was observed (Uesono, Toh-e, and Kikuchi 1997). Ssd1 homologs are important for virulence in a variety of fungal pathogens of both plants and humans (Gank et al. 2008; Tanaka et al. 2007; Thammahong et al. 2019). However, the molecular basis for its role in virulence is not well understood.

*S. cerevisiae* Ssd1 has a mainly cytoplasmic localization, moving to the yeast bud and bud neck during mitosis (Kurischko et al. 2011). This matches the localisation of Cbk1 kinase, which binds and phosphorylates a natively unstructured region of Ssd1, located N-terminal to the RNA binding domains (Fig. 1A) (Gogl et al. 2015; Jansen et al. 2009; Kurischko et al. 2011; Weiss et al. 2002). Transcripts associated with Ssd1 were enriched for mRNAs encoding cell wall biogenesis proteins and Ssd1 was shown to repress their translation (Hogan et al. 2008; Hose et al. 2020; Jansen et al. 2009). An emerging model from these data is that Cbk1 and Ssd1 associate with new buds as yeast cells start to divide. In this model, Ssd1 may help to ensure localized translation at the bud, suppressing translation unless Ssd1 is phosphorylated by bud-localized Cbk1 (Jansen et al. 2009). Consistent with this model, Ssd1 is dispensable, whereas loss of Cbk1 is lethal when wild type Ssd1 is present (Jansen et al. 2009; Kurischko et al. 2005). Loss of Cbk1 results in a strong cell separation phenotype that is suppressed by deletion of *SSD1* (Bidlingmaier et al. 2001; Du and Novick 2002; Racki et al. 2000). The inability to relieve translational repression of cell wall remodeling enzymes may prevent bud growth, leading to cell growth arrest.

**Figure 1.**
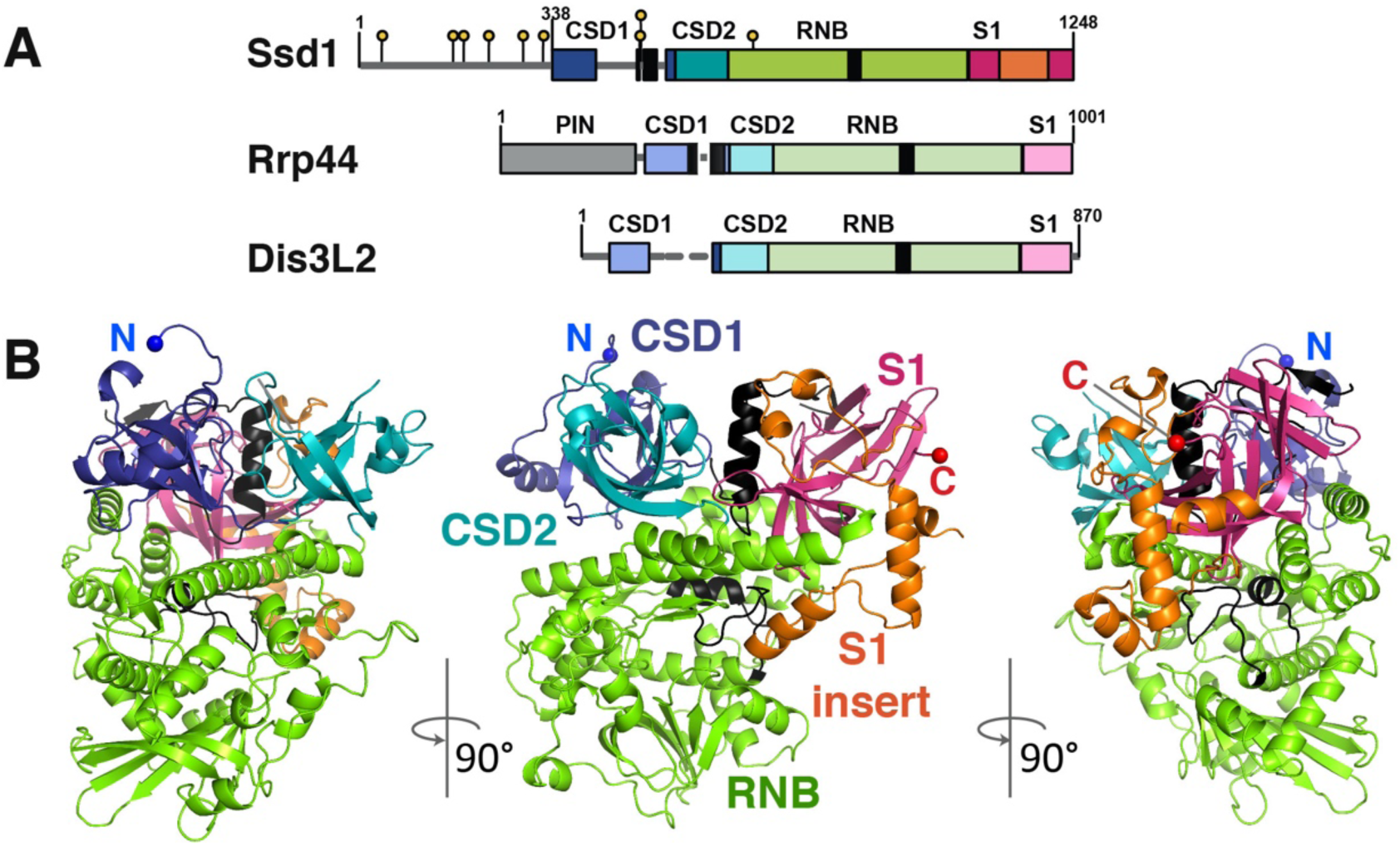
A 1.9 Å crystal structure of Ssd1 reveals conservation of fold with RNase II enzymes. (**A**) Domain overview of Ssd1 and the related proteins Rrp44 (Dis3, numbering is for yeast) and Dis3L2 (numbering is for mouse). Boxes indicate folded domains with separating grey lines indicating natively unstructured regions; yellow lollipops indicate phosphorylation sites of Ssd1. Black boxes equate to features coloured black in the structural figures. (**B**) Structure of Ssd1 observed from three different viewpoints: N-terminal side view, front view and C-terminal side view. The domains are coloured to match those in (**A**).

How Ssd1 recognises RNA and prevents translation is not well understood, mechanistically. Many Ssd1p-associated transcripts have a common C/U-rich sequence motif, termed the Ssd1-enriched element (SEE) (Hogan et al. 2008). However, the exact binding sites of Ssd1 on these RNAs are not known. The SEE element is enriched in 5’ untranslated regions (5’UTRs) of Ssd1-associated transcripts (Hogan et al. 2008), but reporter gene experiments did not clearly identify sequence elements that confer Ssd1-dependent regulation (Wanless, Lin, and Weiss 2014). The SEE sequence element occurs internally in mRNAs, and so is unlike the 3’ terminal elements recognized by RNase II family nucleases, such as Dis3L2 that recognises terminal oligo(U) sequences.

RNase II proteins have a conserved domain structure, consisting of two cold-shock RNA-binding domains, N-terminal to a central catalytic RNB domain, and a C-terminal S1 RNA-binding domain (Fig. 1A)(Frazao et al. 2006; Reis et al. 2013). Here, we present a 1.9 Å X-ray crystal structure of *S. cerevisiae* Ssd1p and show that it retains this domain architecture. The absence of enzymatic activity in Ssd1p arises both from mutation of active site residues and the exploitation of loop elements that likely regulate activity of DIS3-family enzymes. Precise mapping of Ssd1p binding sites transcriptome-wide *in vivo* by UV crosslinking and analysis of cDNAs (CRAC), showed that Ssd1p recognizes a bipartite element than encompasses the previously identified SEE motif, along with a second upstream motif. Finally, the structure gives important insights into the evolution of regulatory motifs within the RNase II enzyme family.

## Results

### Ssd1 retains the domain architecture of DIS3 family nucleases

We aimed to understand the molecular basis for the function of Ssd1 as an RNA binding protein, rather than an enzyme. We solved the crystal structure of the core folded domains of Ssd1. This truncated protein lacks the first 338 residues that are predicted to be natively unstructured (Fig. 1A). Despite the low sequence identity between Ssd1 and yeast Rrp44 or mouse Dis3L2 (22% and 27%, respectively), the structure was solved to 1.9 Å by molecular replacement using fragments of both Rrp44 and Dis3L2.

The overall structure of Ssd1 retains the RNB family domain organization: two N-terminal β-barrel cold shock domains (CSD1 and CSD2) sit at the mouth of a funnel-shaped RNB fold. Opposing CSD1 and CSD2 is a C-terminal β-barrel S1 domain (Fig. 1A,B). Several loop regions (415-484, 492-497, 530-535, 562-578, 1190-1193) could not be assigned in the structure. In total, around 12% of the structure was not visible in the map and could not be built (Fig. S1). The refined model shows good stereochemistry with final *R*_work_ and *R*_free_ of 20.5 % and 22.4%, respectively (Table I).

**Table I.**
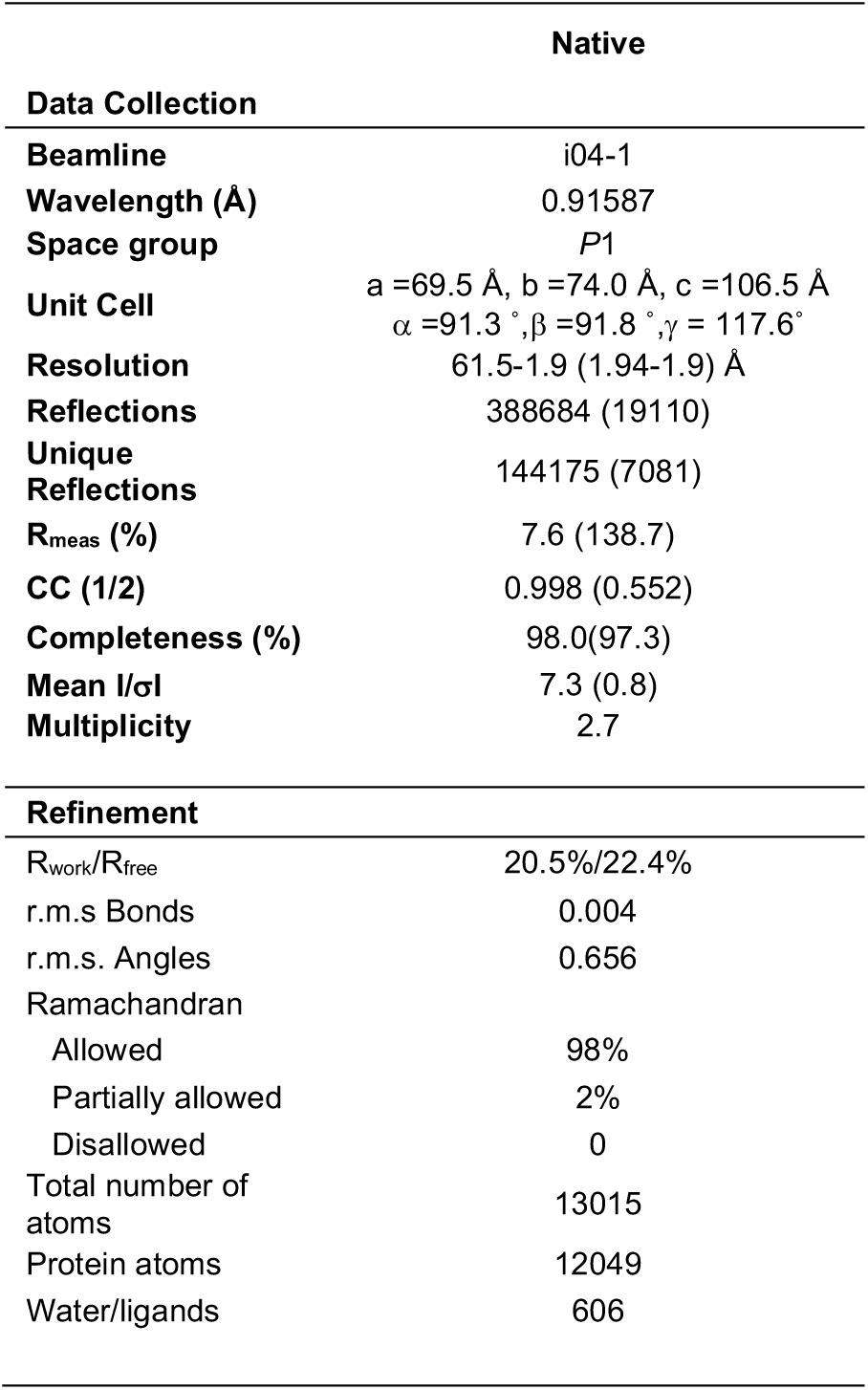

Relative to bacterial RNase II enzymes, Ssd1 contains two insertion elements that are likely to be functionally important. CSD1 is interrupted by an insertion in the loop between strands β4 and β5 (Fig. 1A, Fig. S1), only a portion of which could be assigned in the model (Fig. S2A). Similar insertions are present at the same position in Rrp44 and Dis3L2, so this may be a feature of DIS3 family proteins (Fig. S1A). An additional, Ssd1-specific insertion (residues 1119 to 1204) interrupts the S1 domain (Fig. 1, Fig. S1).

### Structural and sequence changes underlie loss of nuclease activity

Four structural alterations contribute to the loss of nuclease activity in Ssd1. First, active RNase II nucleases have a cluster of four acidic residues that coordinate a divalent cation required for catalysis (Frazao et al. 2006; Zuo et al. 2006). These are absent in Ssd1 (Uesono, Toh-e, and Kikuchi 1997), with the structurally equivalent residues being Ser704, Val709, Glu711 and Phe712; this configuration is unable to coordinate a divalent cation (Fig. 2A).

**Figure 2.**
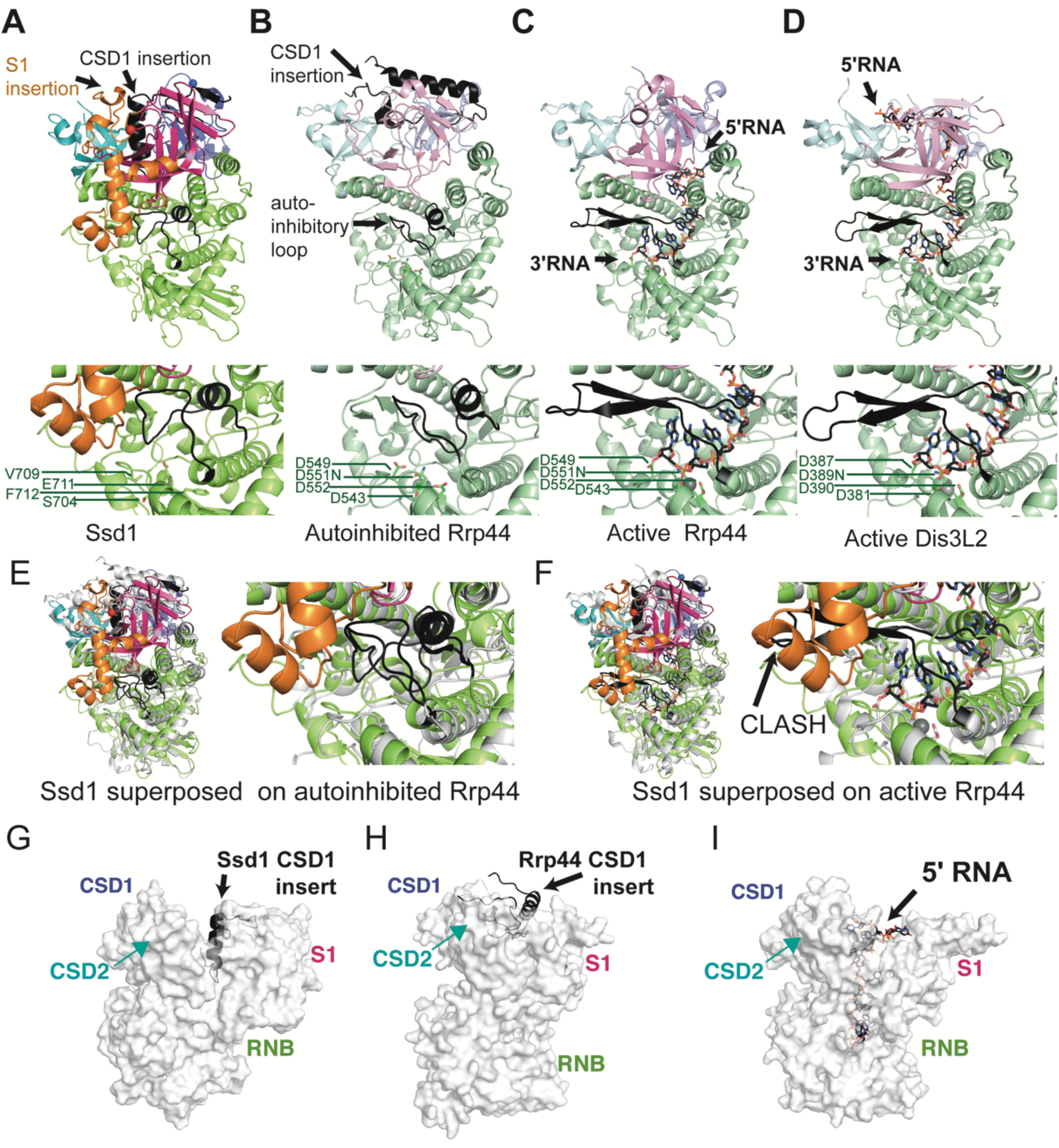
Ssd1-specific structural alterations lock it in an inactive state. Comparison of Ssd1 (**A**) with autoinhibited Rrp44 (**B**) (2wp8), active Rrp44 (**C**) (2vnu) and active Dis3L2 (**D**) (4pmw) using the C-terminal side view. Domains are coloured to match Fig. 1A. The CSD1 insertion and autoinhibitory loop are shown in black. RNA bound to active Rrp44 and Dis3L2 is shown as sticks, with black carbon atoms, and with Mg^2+^ ions shown as grey spheres. Under each structure is a zoomed in view of the equivalent active site residues, shown as sticks. The PIN domain of Rrp44 and associated exosome subunits are omitted for clarity. (**E**) A zoomed view of the active site region of autoinhibited Rrp44 superposed on Ssd1. (**F**) A zoomed view of the active site region of RNA-bound Rrp44 superposed on Ssd1, showing clashes between the reordered autoinhibitory element (black) and the Ssd1-specific S1 insertion (orange). The equivalent segment in Ssd1 is green and occupies the same space as the 3’ end of the RNA. (**G**) Insertion of the CSD1 insert (black) into the funnel region between the CSD and S1 domains of Ssd1 (grey surface). The view is the “front” view from Fig. 1A. (**H**) Insertion of the CSD1-insert (black) into the funnel region of autoinhibited Rrp44 (grey surface). (**I**) Similar view of Dis3L2 compared to Ssd1 and Rrp44 in (**G**) and (**H**), showing the path of the RNA through the funnel.

Of the available structures of eukaryotic RNase II nucleases, Ssd1 most closely resembles the conformation of Rrp44 when in a complex with two proteins of the exosome complex (Bonneau et al. 2009)(compare Fig. 2A, 2B). In this structure, a loop segment within the RNB domain of Rrp44 forms an α-helix, which blocks the channel that is occupied by the RNA substrate during catalysis (Fig. 2B). A similar configuration is observed in Ssd1, representing a second structural change that contributes to loss of activity (Fig. 2A, Fig. S2B). This segment blocks the lower tract of the active site. In contrast, in structures of Rrp44 and Dis3L2 engaged with RNA substrates this loop is rearranged to form a β hairpin motif outside of the active site, allowing RNA substrates to be accommodated (Fig. 2C, 2D)(Faehnle, Walleshauser, and Joshua-Tor 2014; Lorentzen et al. 2008). Indeed, this active conformation is also observed in other RNase II nucleases such as Dss1 and *E. coli* RNAse II (Fig. S3A, S3B)(Frazao et al. 2006; Razew et al. 2018).

In Rrp44, the mobile autoinhibitory element can switch between active and inactive states (Fig. 2B). In contrast, this segment in Ssd1 is fixed in place by a third structural change: the insertion segment in the S1 domain that is apparently unique to Ssd1 (Fig. 2A, 2E, orange segment). The S1 insertion packs against the RNB domain and stabilizes the inhibitory conformation of the loop (Fig. 2E). Superposition of Ssd1 on active Rrp44 shows that the S1 insertion would have a steric clash with the autoinhibitory segment in the active configuration (Fig. 2F). We conclude that the S1 insertion locks the autoinhibitory segment in place, ensuring a fixed, inactive conformation.

A fourth structural element prevents access of RNA to the former active site of Ssd1. Active enzymes, such as Dis3L2 (Fig. 2D), bacterial RNase II (Fig. S3B), and human DIS3 (Fig. S3C), anchor substrate RNA at the mouth of the funnel created by CSD1, CSD2 and S1 domains, allowing the RNA to thread into the active site (Faehnle, Walleshauser, and Joshua-Tor 2014; Frazao et al. 2006; Weick et al. 2018). The CSD1 insertion in Ssd1, which is only partially assigned in this structure, folds into an α-helix that blocks the mouth of the funnel, excluding RNA binding at this surface (Fig. 2G). The S1 insertion element also packs against the CSD1 insertion, further stabilising this conformation (Fig. 1B, 2A). A CSD1 insertion at an equivalent position is not present in bacterial RNase II but is present in Dis3L2, human DIS3 and yeast Rrp44. In Dis3L2, the insertion was engineered out of the construct used for crystallization and so cannot be observed (Faehnle, Walleshauser, and Joshua-Tor 2014). A large portion of the Rrp44 CSD1 insertion is ordered in the autoinhibited structure of yeast Rrp44 and, similar to Ssd1, blocks the upper cavity (Bonneau et al. 2009) (Fig. 2H). In Dis3L2 and other RNase II enzymes, this cavity is required for RNA access (Fig. 2I, Fig. S3B, S3C). It should be noted, however, that RNA substrates typically access yeast Rrp44 active site by tunnelling between CSD1 and the RNB domain (Fig. 2C) while the human homologue DIS3 has been observed to bind RNA in a similar mode to Dis3L2 (Fig. S3C) (Makino et al. 2015; Weick et al. 2018). The structural elements that block RNA access to the central channel of Ssd1 may have evolved from regulatory switches in ancestral proteins that have become fixed in the “off” state.

### Ssd1 uses an alternative surface for RNA binding

As the insertion elements in Ssd1 block the central funnel of the RNAse II fold, Ssd1 cannot bind RNA using the same mode as its enzyme relatives. We examined the surface properties of the protein to search for alternative RNA binding sites. Our previous evolutionary analysis of fungal Ssd1 and Dis3L2 homologues places Ssd1 as the sole *S. cerevisiae* homologue of Dis3L2, and shows that the CSDs are highly conserved in Ssd1 homologues in ascomycete and basidiomycete fungi (Ballou, Cook, and Wallace 2020). Based on this analysis, we segregated high confidence Ssd1 sequences from other Dis3L2 homologues that retain negatively charged residues at the active site, i.e. we excluded sequences that are likely to be active nucleases. Using a multiple sequence alignment consisting of 91 high confidence Ssd1 homologues across fungi (Supplementary File 1), surface conservation was calculated using the CONSURF server (Ashkenazy et al. 2016; Landau et al. 2005). This revealed an extensive conserved surface around the two CSD domains and the RNB domain (Fig. 3A). The conserved area coincides with a large surface patch of positive charge, consistent with binding to nucleic acids (Fig. 3B). This candidate RNA-binding surface provides further evidence that RNA-binding by Ssd1 is distinct from substrate recognition by Rrp44 and Dis3L2 enzymes.

**Figure 3.**
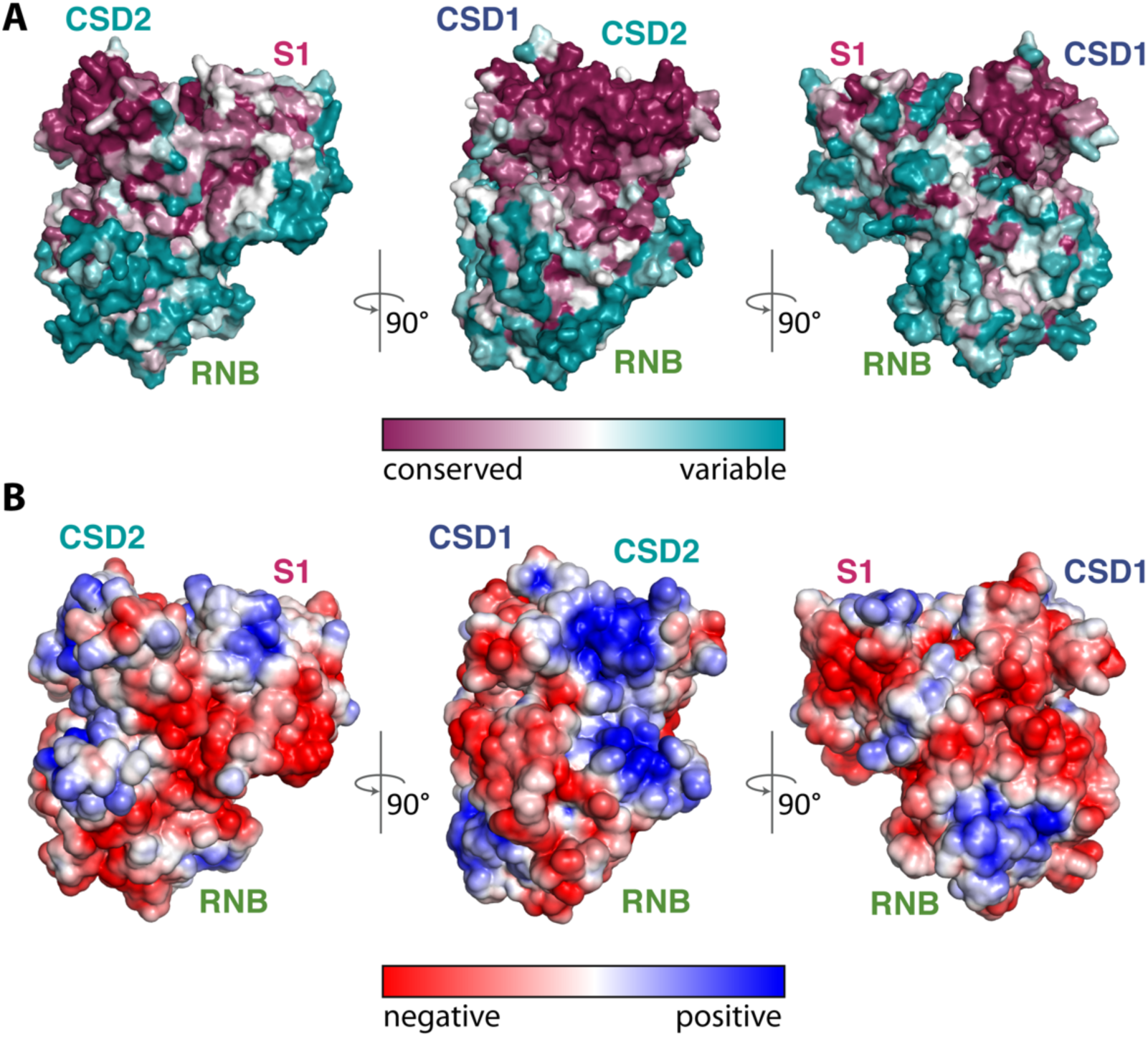
A conserved positively charged surface is a candidate RNA-binding site. (**A**) Ssd1 shown as a van der Waals surface coloured by conservation showing front, side and back views. Front and side views match those in Fig. 1. Conservations scores were calculated using CONSURF. (**B**) The same views of Ssd1 showing surface electrostatics on a solvent accessible surface, calculated using APBS. The gradient is from −3 to +3 kT/e.

### Ssd1 associates primarily with 5’UTRs of mRNAs encoding cell wall proteins

To locate the precise RNA binding sites of Ssd1, we carried out UV cross-linking and analysis of cDNAs (CRAC) (Granneman et al. 2009) on a yeast strain where Ssd1 was expressed with a C-terminal HTP (hexahistidine-TEV cleavage site-Protein A) tag from the endogenous locus. To determine whether affinity tags at the N- or C-terminus of Ssd1 would affect its *in vivo* activity, we used two functional assays for which *ssd1* phenotypes are well characterized. Calcofluor white (CFW) binds to chitin in fungal cell walls, and *ssd1Δ* strains are sensitive to CFW concentrations in the range of 10 to 100µM (Kaeberlein and Guarente 2002). Neither N-terminal nor C-terminal tags on endogenous Ssd1 increased sensitivity of cells to CFW, whereas *ssd1Δ* or a Ssd1 truncation that lacks the first 338 residues (ΔN338 Ssd1, equivalent to the construct used for structural studies) were both highly sensitive to treatment (Fig. 4A). Loss of Ssd1 also reduces “induced thermotolerance” in yeast, where a mild heat shock protects cells from death in subsequent severe heat shock (Mir, Fiedler, and Cashikar 2009). Again, neither N-terminal nor C-terminal tags on endogenous Ssd1 altered their thermal tolerance. Interestingly, ΔN338 Ssd1 showed a thermal tolerance phenotype similar to wild type rather than to *ssd1Δ* (Fig. 4B).

**Figure 4.**
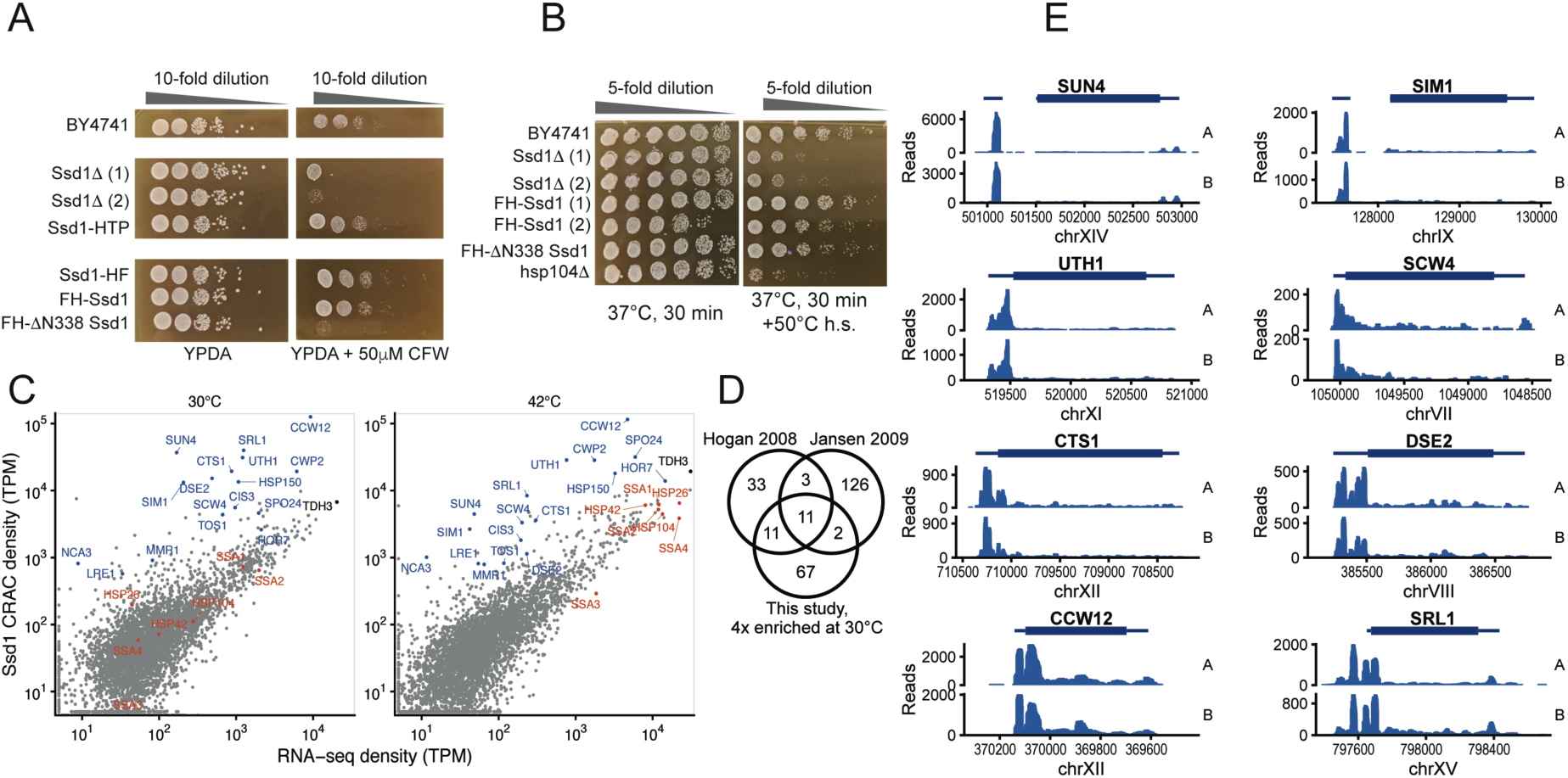
Ssd1 binds to 5’UTRs of its target transcripts *in vivo*. (**A**) Wild type and Ssd1 mutant yeast strains grown at 30°C on YPDA without or with 50µM calcofluor white (CFW). (**B**) Wild type and Ssd1 or Hsp104 mutant strains grown overnight at 30°C and then incubated at either 37°C for 30 mins or using an induced thermotolerance protocol (37°C for 30 mins followed by heat shock at 50°C for 30 mins before plating and growth at 30°C). (**C**) Ssd1-bound CRAC read density compared to RNA-seq reads in transcripts per million (TPM, mean over two biological replicates), aligned to full-length transcripts including annotated UTRs. Selected Ssd1 targets are highlighted in blue and selected heat-induced transcripts in red. (**D**) Comparison of Ssd1-bound mRNAs reported by CRAC analysis with previous RNA immunoprecipitation and microarray studies, that were also conducted in rich media at 30°C. We conservatively report transcripts that are 4-fold enriched in Ssd1 CRAC reads compared to RNA-seq, and with at least 20 TPM in the RNA-seq data. (**E**) Unnormalized CRAC read counts (pileups) on selected Ssd1-bound transcripts from two biological replicates at 30°C, aligned to the genome, with 5’UTRs oriented on the left.

CRAC studies were done in biological duplicates on cells growing exponentially in synthetic medium at 30°C, and following heat shock at 42°C for 16 minutes, a condition in which total Ssd1 binding to RNA markedly increases (Bresson 2020). Expression of tagged constructs was verified by western blot (Fig. S4A); tandem immunoprecipitation of these samples efficiently recovered cross-linked RNA (Fig. S4B), while a negative control did not. CRAC data derived from these samples were reproducible, whereas a negative control strain with no tag gave low read counts and low correlation with the Ssd1-HTP results (Fig. S4C). We compared the CRAC reads to poly(A)-enriched RNA-seq from the background (untagged) yeast strain grown in matched conditions (Bresson 2020), which were also reproducible (Fig S4D). Counting reads on full-length transcripts, and normalising by transcript length and total density to transcripts per million (TPM)(Wagner, Kin, and Lynch 2012), revealed a clear enrichment for a small proportion of mRNAs (Fig 4C). Many of the identified Ssd1-bound mRNAs encode proteins required for cell wall biogenesis or septum remodelling, including *SUN4, SIM1, UTH1, SCW4, CTS1, DSE2, CCW12*, and *SRL1* (Fig. 4C, 4D), in agreement with previous studies (Hogan et al. 2008; Hose et al. 2020; Jansen et al. 2009). These mRNAs are enriched for Ssd1 binding regardless of heat shock, despite dramatic changes in RNA expression levels and an increase in overall Ssd1 binding between conditions (Bresson 2020). By contrast, mRNAs encoding heat-shock proteins are increased in their expression levels on heat shock by several orders of magnitude. However, they are not enriched in Ssd1-binding when the increase in their mRNA abundance is taken into account (Fig 4C).

We next looked at the profile of Ssd1-bound reads within individual transcripts (Fig 4E), finding that Ssd1 is overwhelmingly and reproducibly bound to 5’UTRs of its target transcripts. For the paralogous genes SUN4 and SIM1, Ssd1 reads are concentrated in exon 1, upstream of a 5’UTR intron. Some targets have a series of distinct peaks in the 5’UTR, in some cases extending into the coding sequence (CCW12, CTS1, SRL1) (Fig 4E). Additional, smaller peaks in the 3’UTR were also observed in some cases (SCW4, SRL1). These data show Ssd1 to be targeted to discrete regions of specific transcripts *in vivo*; 5’UTR binding is consistent with the reported role of Ssd1 as a repressor of translation.

### MEME analysis reveals three sequence motifs associated with Ssd1 cross-link sites

We next investigated the sequence determinants of Ssd1’s binding specificity. Using the sequences found in CRAC peaks (FDR < 0.05), MEME analysis was carried out to find sequence motifs enriched on Ssd1 targets. Consistent with previous reports, MEME analysis of the top 100 Ssd1-associated peaks identified 59 occurrences of a general motif CNYUCNYU, similar to the previously reported SEE motif AKUCAUUCCUU (Fig. S5A) (Hogan et al. 2008; Wanless, Lin, and Weiss 2014). Notably, transcripts with high confidence Ssd1 binding sites generally have more than one CNYUCNYU motif within the 5’UTR, including most of the transcripts shown in Fig. 4D. For example, UTH1 and SRL1 each have 5 CNYUCNYU sites in their 5’UTRs (Fig. S5B).

CRAC allows precise mapping of cross-linked sites because the RNA-protein cross-link leaves a moiety on the RNA after protease digestion. This “cross-linking scar” can cause Superscript family reverse transcriptase to skip bases, which appear as deletions in the aligned sequences. Notably, across the dataset, a peak of higher frequency deletions 2-4 nucleotides upstream of the CNYUCNYU motif was observed (Fig. S5B). This is exemplified by the *SUN4* 5’UTR (Fig. 5A) where two nearby CNYUCNYU sites both have a high proportion of deletions a few bases upstream of the motif. This confirms that the motifs are in contact with Ssd1 *in vivo*. These deletions can be mapped only to a 4nt region, as they are ambiguous substitutions of CUCU to CU and UUUU to UU.

**Figure 5.**
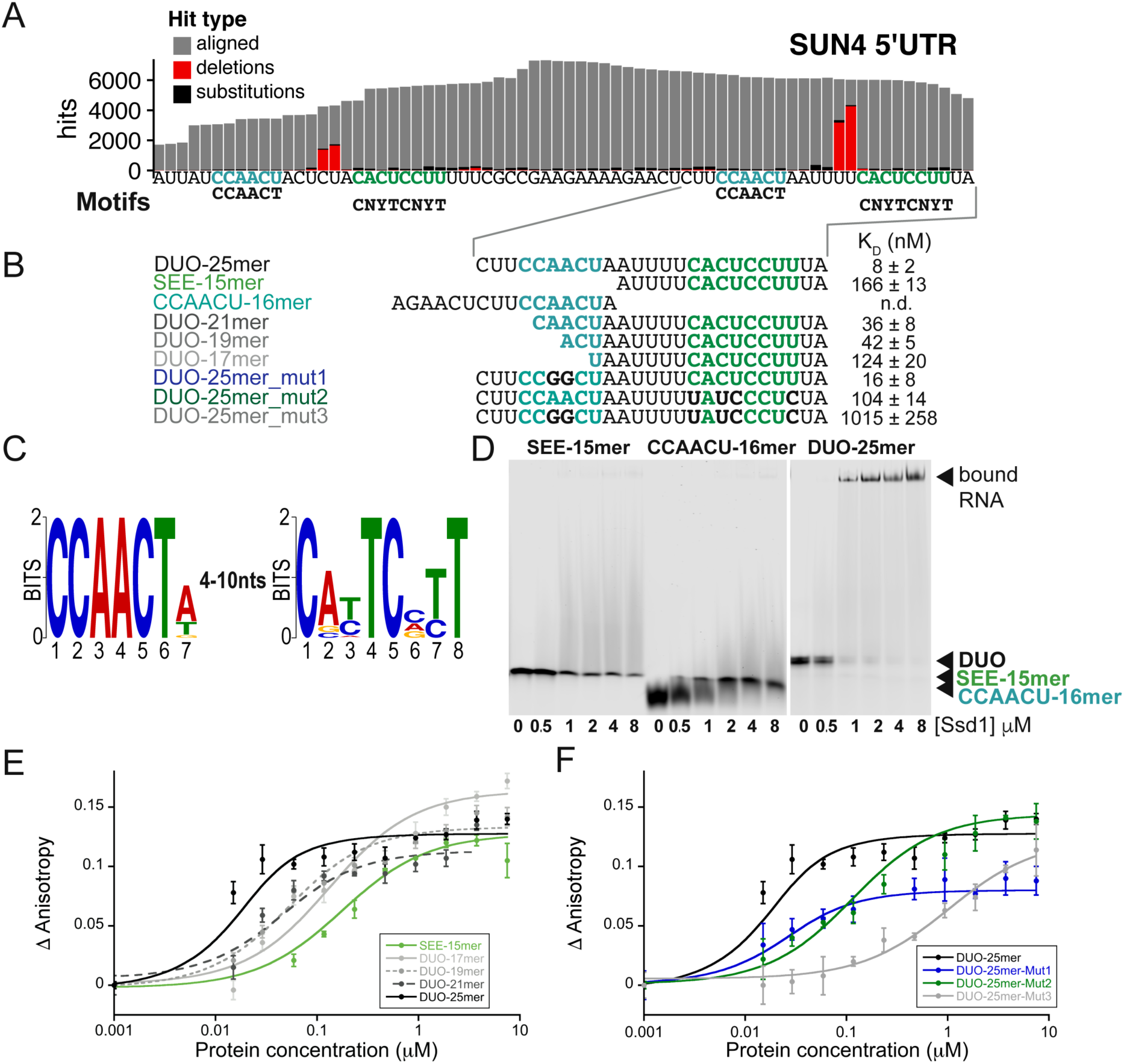
Ssd1 binds directly to a bipartite motif found in target 5’UTRs. (**A**) Zoomed in view of CRAC reads on SUN4 5’UTR with CCAACU and CNYUCNYU motif positions, showing mutations and deletions indicative of RNA-protein crosslinking sites. (**B**) Overview of oligomer sequences used in EMSA and fluorescence anisotropy binding assays with calculated K_D_ values in nM. (**C**) Sequence logo (in DNA alphabet) of two Ssd1-enriched motifs found by MEME analysis of the 100 top peaks in CRAC read data (**D**). EMSA binding assays for SUN4 5’UTR oligonucleotides. RNA probes were present at 0.5 µM (**E**) Fluorescence anisotropy data, with fitted curves, used to calculate K_D_ values of different lengths of RNA derived from the SUN4 5’UTR. (**F**) Fluorescence anisotropy data, with fitted curves, used to calculate K_D_ values of RNA derived from the SUN4 5’UTR with mutations.

We further noted that the two Ssd1-associated copies of the CNYUCNYU motif in the SUN4 5’UTR are preceded by a CCAACU motif (Fig. 5A, B). Moreover, the Ssd1 cross-linking sites lie between the CNYUCNYU motif and the CCAACU motif. By contrast, a third copy of the CNYUCNYU motif in the SUN4 5’UTR does not have this upstream motif, and has far fewer Ssd1 CRAC reads (Fig. S5B). The MEME analysis found a CCAACUV motif weakly enriched across the dataset, invariably appearing 1-4nt upstream of CNYUCNYU peaks (Fig 5C, S5A). This indicates that the combination of these two motifs, as seen in *SUN4*, is a commonality among Ssd1 targets. In addition to these two motifs, a significantly enriched purine-rich motif was also observed, which we do not pursue further (Fig. S5A).

### Short sequence motifs are not sufficient for binding to Ssd1

To determine whether the CNYUCNYU motif is sufficient to bind to Ssd1, we carried out electrophoretic mobility shift assays (EMSA) with a native sequence corresponding to one of the tandem CNYUCNYU motifs of *SUN4* 5’UTR and ΔN338 Ssd1 (Fig. 5A,B,D). However, binding of Ssd1 to this RNA oligomer was barely detectable (Fig. 5D). As the MEME analysis indicated that the CCAACU motifs is also enriched in several Ssd1 targets, we tested the CCAACU motif in our EMSA assay but saw no significant binding (Fig. 5D). However, when we carried out the same assay using a longer “DUO” RNA that encompasses both motifs, we saw strong production of a specifically shifted band (Fig. 5D). This indicated that either a longer RNA, or the combination of the two sequence elements, or both, are required for efficient Ssd1 binding.

To better understand how these two motifs affect RNA recognition by Ssd1, we used fluorescence anisotropy to measure the binding affinity of Ssd1 to fluorescently-labelled RNA “DUO” oligos that encompassed both motifs, or were progressively shortened from the 5’ end to disrupt the CCAACU sequence (Fig. 5B,E, Fig. S6). A 25mer oligomer encompassing both motifs bound to Ssd1 with a K_D_ of 8 nM, while a 15mer oligomer that encompasses only the CNYUCNYU motif had a K_D_ of 166 nM, consistent with the EMSA data (Fig. 5D,E, Fig. S6). Intermediate sized oligomers of 21, 19 and 17 nt showed progressively weaker binding (Fig. 5B,E, Fig. S6). However, the largest changes in affinity were between the 25mer and 21mer oligomers (8 nM and 36 nM respectively) and between 19mer and 17mer oligomers (42 nM and 124 nM, respectively) (Fig. 5B,E, Fig. S6).

The loss of affinity when RNAs were shortened from the 5’ end suggested that RNA length is important. However, the sequence may also contribute to binding. To further test this, we measured binding affinities for three different mutated oligos. For the first mutant, we altered the central AA bases of the CCAACU motif to GG, to maintain purines at this site but change the base. This alteration had a mild (2-fold) reduction in affinity, in a similar range observed for the DUO-19mer and DUO-17mer oligomers that have a partly truncated CCAACU motif (Fig. 5B,F, Fig. S6). In a second mutation, we switched four of the conserved pyrimidine bases (C-to-U, or U-to-C mutations) of the CNYUCNYU motif. This mutation substantially altered the binding affinity from the low nanomolar range to a K_D_ of 104 nM (Fig. 5B,F, Fig. S6). A third mutation combined the alteration to the CNYUCNYU motif with that of the AA-to-GG alteration in the CCAACU motif. This mutation led to a further ∼10-fold loss of affinity. We conclude that the combined CCAACU and CNYUCNYU motif sequences are important for high affinity binding of Ssd1. Together, these data are consistent with the CCAACU motif contributing sequence specificity in addition to the effect of RNA length.

### Ssd1 binding motifs are conserved across fungi

The Ssd1 binding site is highly conserved in homologous transcripts: we focused on the SUN4 5’UTR by aligning the upstream regions of SUN4 homologs from 7 sequenced species of *Saccharomyces* sensu stricto (Fig. 6A). This sequence logo shows that both the CCAACU and CNYUCNYU motifs are perfectly conserved in the tandem binding site, while nearby sites are more variable. This tandem binding site was previously identified as highly conserved using a phylogenetic hidden Markov model, PhastCons (Siepel et al. 2005). This confirms that specific nucleotides within these two motifs are conserved over 20 million years of evolution in the *Saccharomyces* genus.

**Figure 6.**
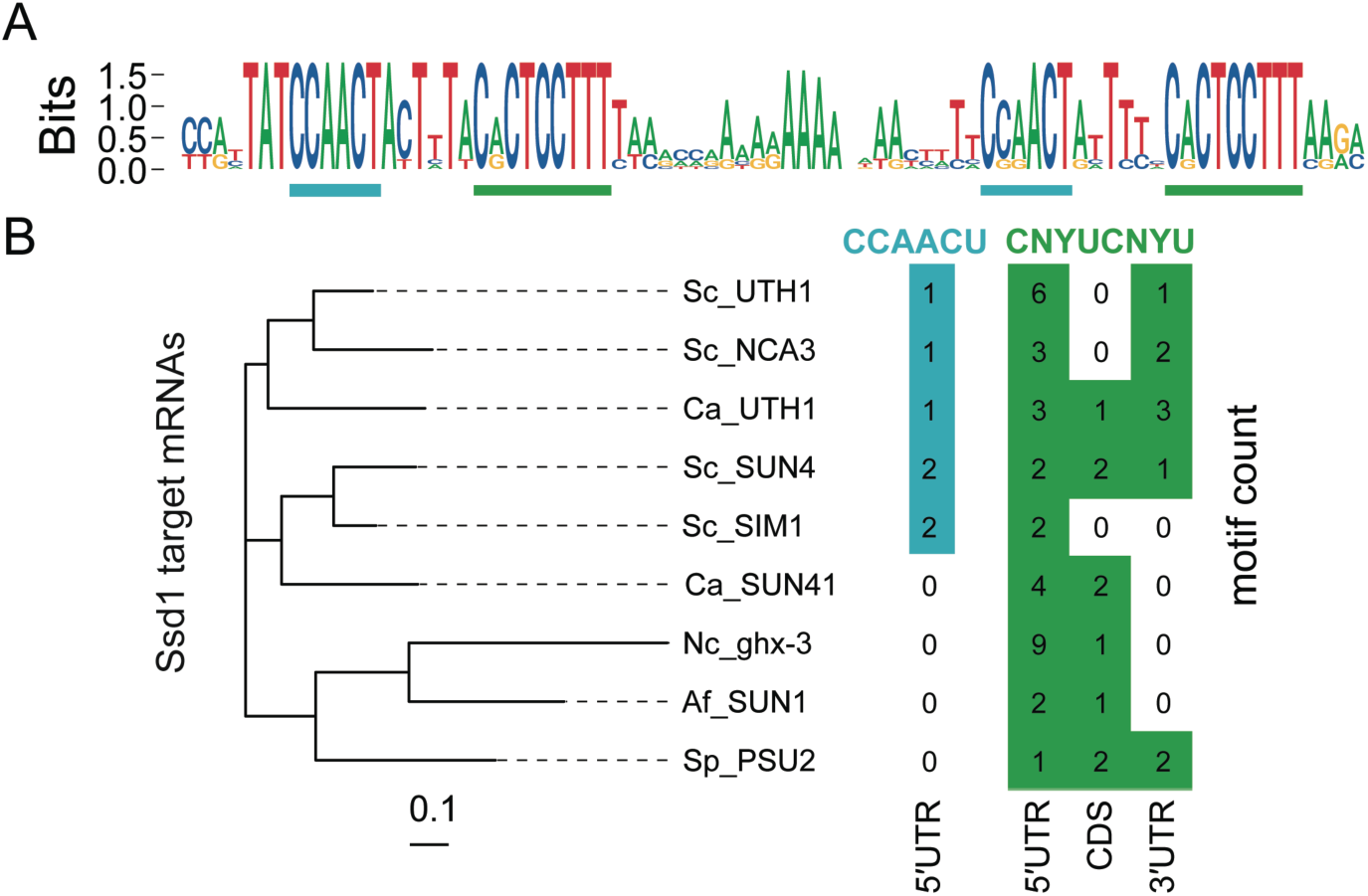
Ssd1 target sites are conserved across fungi. (**A**) Sequence logo (in DNA alphabet) of Ssd1 tandem binding site on SUN4 5’UTR as aligned from 7 species of *Saccharomyces* spp. CCAACU and CNYUCNYU motifs are highlighted as in Fig. 5A. (**B**) Motif counts of CCAACU and CNYUCNYU within transcripts of SUN4 homologues in select ascomycete fungi, aligned to the protein phylogenetic tree on the left. Genes are *S. cerevisiae* SUN4, SIM1, UTH1, and NCA3; *C. albicans* UTH1/SIM1 and SUN41; *N. crassa* ghx-3; *A. fumigatus* SUN1; and *S. pombe* PSU2.

The binding motif of Ssd1 is also conserved at longer evolutionary distances. Indeed, a sequence similar to the SEE motif was reported to be found in 5’UTRs of *Schizosaccharomyces pombe* Sts5, a homologue of Ssd1 that is also a pseudonuclease (Ballou, Cook, and Wallace 2020; Nunez et al. 2016). We focused our search on homologues of SUN4; SUN4 and SIM1 are post-whole genome duplication paralogues, as are UTH1 and NCA3 (Byrne and Wolfe 2005). These putative glucanases are secreted proteins that localise to the bud scar in *S. cerevisiae* (Kuznetsov et al. 2013; Kuznetsov, Vachova, and Palkova 2016). We analysed transcript annotations of SUN4 homologues from other ascomycete fungi *Candida albicans, Neurospora crassa, Aspergillus fumigatus*, and *S. pombe*. We counted the instances of CCAACU in the 5’UTR, and CNYUCNYU in the 5’UTR, CDS, and 3’UTR and compared these instances with a phylogenetic tree of the proteins (Fig. 6B). All of these ascomycete SUN-family genes have multiple CNYUCNYU motifs in the 5’UTR or near to the start codon. For example, *S. pombe* PSU2 has one CNYUCNYU motif in the 5’UTR and two CNYUCNYU motifs in the CDS, respectively 14nt and 38nt downstream of the start codon. We find CCAACU motifs upstream of CNYUCNYU motifs in 5’UTRs only from *S. cerevisiae* and *C. albicans* homologs, suggesting that the bipartite RNA binding motif is particular to the budding yeast clade.

These results indicate that regulation of downstream targets of Ssd1 is conserved over more than 500 million years of ascomycete evolution, in addition to conservation of the upstream regulation of Ssd1 by Cbk1 (Ballou, Cook, and Wallace 2020; Jansen et al. 2009; Nunez et al. 2016).

## Discussion

The Ssd1 structure reveals an inactive pseudonuclease, in which the ancestral path of RNA into the funnel of the active site is blocked by fixed structural elements. These fixed elements are informative about the evolutionary history of this family of proteins. Bacterial RNase II proteins do not appear to have regulatory elements such as insertion of the cold shock domain CSD1 (Fig. 7A). Moreover, the autoinhibitory loop is in an open conformation (Fig. S3B), as observed in many structures of related nucleases, particularly when RNA substrates are present (Fig. 2C, 2D, S3A, S3C). In contrast, the closed, autoinhibited conformation was previously observed only for Rrp44 (Bonneau et al. 2009). The combination of the autoinhibitory segment and the CSD1 insertion, which is able to block the top of the funnel (at least in Ssd1 and Rrp44), suggest that ancestral Dis3-family enzymes may have acquired these two mobile elements to facilitate regulation, by switching between a closed, autoinhibited form and an open, active form (Fig. 7B). In Ssd1 homologs, these segments have been trapped in the “off” state by the Ssd1-specific S1 insertion that packs against both the CSD1 insertion and the autoinhibitory loop (Fig. 7C).

**Figure 7.**
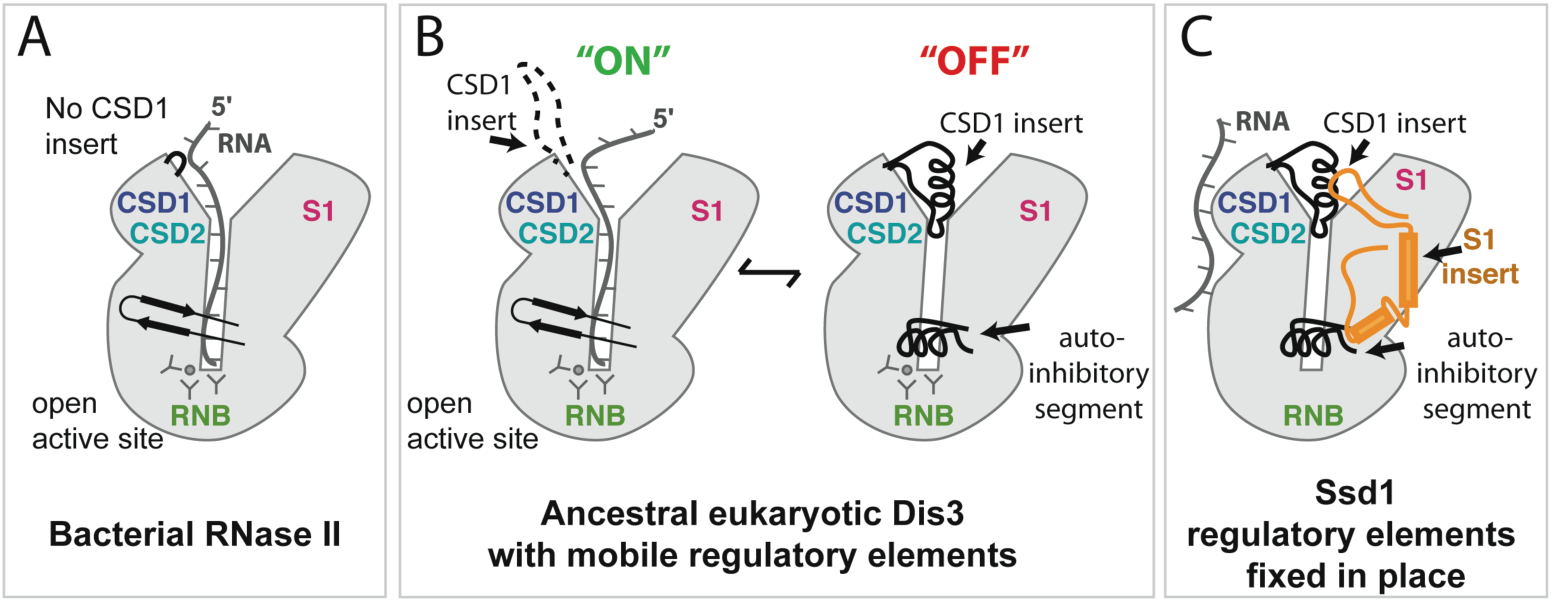
Evolution of RNase II enzymes and pseudoenzymes. (**A**) Bacterial RNase II has a domain structure that is conserved in evolution but lacks the eukaryotic-specific insertions. RNA accesses the active site by funnelling into the core of the protein. (**B**) An ancestral Dis3/Dis3L2 enzyme may have acquired mobile regulatory elements that allow the enzyme to be finely regulated. The “ON” state resembles that of the bacterial enzyme while the “OFF” state uses the CSD1 insertion and the autoinhibitory segment to block the funnel. (**C**) The autoinhibitory elements have been fixed in place in Ssd1 by the S1 insertion element and the active site residues have been lost. A new RNA binding site has been acquired.

An important consequence of Ssd1 having acquired a permanent “off” state is that the funnel-shaped RNA binding site that recognises RNA 3’ ends is blocked. Instead, Ssd1 has gained a new RNA binding site that allows it to bind internally on transcripts. It is likely that the new RNA binding site is a conserved, positively-charged region on the outer face of the two CSDs (Fig. 7C). Our previous evolutionary analysis indicated that Ssd1 is the closest yeast homologue of Dis3L2 and that loss of nuclease function in Dis3L2 homologues has occurred independently in multiple fungal lineages (Ballou, Cook, and Wallace 2020). These analyses also indicated that the CSDs are the most highly conserved part of Ssd1. We speculate that an Ssd1 ancestor was a bifunctional RNA degrading nuclease and RNA binding protein, and that the latter function has been preserved in preference to the nuclease activity.

RNA cross-linking *in vivo* confirmed that Ssd1 binds to 5’UTRs of transcripts encoding a subset of cell wall proteins. This provides further evidence for a distinct mode of RNA binding, because these Ssd1 binding sites are internal to RNAs while Rrp44 and Dis3L2 bind to the terminal 3’ ends of RNA. Ssd1-bound transcripts contain the previously identified, conserved CNYUCNYU motif. However, this motif is not sufficient for tight Ssd1 binding *in vitro*. We demonstrate that a longer bipartite motif, encompassing an upstream CCAACU sequence, in addition to the CNYUCNYU motif, is required for Ssd1 to bind short stretches of RNA *in vitro* in the nanomolar range. We also observe that high affinity Ssd1 transcripts have more than one binding site for the protein. Moreover, homologues of these Ssd1-bound transcripts in other ascomycete fungi also have multiple CNYUCNYU binding motifs, mostly in their 5’UTRs. These observations suggest that Ssd1 binds cooperatively on RNA when more than one copy of the CNYUCNYU motif is present but the CCAACU motif absent. It is also possible that additional cis-elements and additional RNA binding proteins might contribute to stabilisation of Ssd1 on target transcripts.

Ssd1 has been proposed to act as a translational repressor and we speculate that tight (low nM) binding to specific sequences in 5’UTRs can physically block scanning of translation pre-initiation complexes. Regardless of the mechanism, translational repression is likely to be conserved among fungi that encode Ssd1 homologues. Given that Ssd1 is a virulence factor for many fungal pathogens, understanding both the molecular mechanism and the cellular functions of this translational block are important goals for future work.

## Materials and methods

### Expression and purification

The N-terminal deletion construct of Ssd1 Δ338 was cloned as a His-tagged fusion protein into a pET-based expression vector (Supplementary File 2). The protein was expressed in the *E. coli* strain BL21-codon plus-RIL (*DE3*) and was grown in 2XTY media. Cultures were induced with 0.3 mM isopropyl-B-D-1 thiogalactopyranoside (IPTG) overnight at 20°C. Cells were lysed using a cell disruptor (Constant Systems) in lysis buffer (20 mM Tris-HCl pH 8.0, 200 mM NaCl, 10 mM Imidazole, 1 mM β-mercaptoethanol) in the presence of protease inhibitor cocktail (Roche) and DNAse I (Sigma Aldrich). The clarified lysate was bound to Ni-NTA resin (Sigma Aldrich) in batches. The Ni-NTA resin was packed in a XK16/60 column and the unbound protein was washed out using the lysis buffer. The bound protein was eluted with 20 mM Tris-HCl pH 8.0, 200 mM NaCl, 500mM Imidazole, 1mM β-mercaptoethanol. The protein was dialyzed in 20 mM Tris-HCl pH 7.5, 100 mM NaCl, 1 mM dithiothreitol (DTT) in the presence of 3C-protease to cleave off the His-tag. The protein was further purified from nucleic acids by passing through a heparin sepharose column and eluted using 20 mM Tris-HCl pH 7.5, 1000 mM NaCl, 1 mM DTT in a salt gradient. The pure protein was finally purified by size exclusion chromatography (Sephadex 200 column (GE healthcare)) in 20 mM HEPES pH 7.5, 150 mM NaCl, 1 mM DTT.

### Crystallization and structure solution

Ssd1 Δ338 was concentrated to 11.5 mg/ml and crystallized in sitting drops containing a well solution of 50 mM Tris-HCl pH 8.0, 25% PEG 400 at RT. Crystals were cryoprotected in 50 mM Tris-HCl pH 8.0, 30% PEG 400 and flash cooled in liquid nitrogen. Initial crystals diffracted to 3.9 Å. Crystal diffraction quality was improved after reducing the protein concentration in the sitting drops to 9.4 mg/ml. Data were collected at Diamond Light Source (DLS) on beamline i04-1. Data from crystals diffracting to 1.9 Å, with space group *P*1, were obtained and indexed and reduced using the automated data processing suite at DLS (Winter 2010). The structure was solved by molecular replacement by using separate domains from Rrp44 (PDBid 2vnu (Lorentzen et al. 2008)) and DIS3L2 (PDBid 4pmw (Faehnle, Walleshauser, and Joshua-Tor 2014)) with PHASER (McCoy et al. 2007) in MR mode. Two molecules were found in the asymmetric unit. Sub-fragments of the structure of yeast Rrp44 (Dis3) were used as search models and RNB domains were placed first followed by the two N-terminal CSDs from Rrp44 as a single search model. The S1 domain from DIS3L2 was placed last. After initial placement of these sub-fragments, the model was refined using MORPHMODEL in PHENIX (Liebschner et al. 2019; Terwilliger et al. 2012), followed by rounds of rebuilding in COOT (Emsley and Cowtan 2004) and refinement in PHENIX. The final model was assessed for quality using MOLPROBITY (Chen et al. 2010). Figures were prepared with IBS (Liu et al. 2015) and pymol (Schrodinger 2015).

### RNA preparation

All RNA oligonucleotides were synthesized by Biomers GmbH and reconstituted in H_2_O to a final concentration of 1mM (Supplementary File 2). For EMSA assays and fluorescence anisotropy, RNA oligomers were labelled during synthesis at the 5’ end with fluorescent dyes, DY681 and Cyanine3, respectively.

### Electrophoretic mobility shift assays

Binding reactions, containing 0.5 µM RNA 5’-labelled with fluorescent dye DY681 and increasing concentrations of Ssd1 protein, were incubated on ice in binding buffer (20mM HEPES pH 7.5, 150mM potassium acetate and 5mM magnesium acetate). After 1 hour of binding, all samples were mixed with native gel loading buffer containing 0.25% bromophenol blue, 0.25% xylene cyanol, 50% glycerol and 4 µl was loaded onto an 8% native polyacrylamide gel. After 1.5 hour or 3 hours at 2W, the gel was scanned on a LICOR Odyssey fluorescent infrared scanner at 700 nm. Images were converted to greyscale using LICOR Image Studio Software.

### Fluorescence anisotropy

Fluorescence anisotropy assays were carried out in a final volume of 100 μL in black, 96-well plates using a SpectraMax M5 multimode plate reader (Molecular Devices). A total of 20 nM Cy3-labelled ssRNA (in 20 mM HEPES pH 7.5, 150 mM NaCl, 1mM DTT) was incubated with increasing concentrations of Ssd1ΔN338 protein for 15 and 30 min on ice. Anisotropy was measured using 530 and 565 nm wavelengths for excitation and emission, respectively. Experimentally obtained anisotropy was plotted against protein concentration to determine the equilibrium dissociation constant, K_D_, for binding of the labelled RNA oligomer to protein (KaleidaGraph V4.5.4 Synergy Software). The binding curves are described by the following equation:

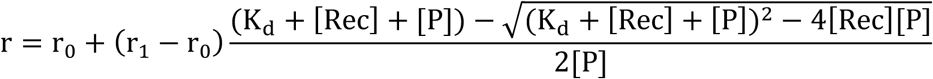

where r is the observed anisotropy, r_0_ is the anisotropy of free Cy3-labelled RNA, r_1_ is the anisotropy of fully bound RNA, [Rec] is the protein concentration, [P] is the Cy3-labelled RNA concentration and K_D_ is the dissociation constant for the interaction. Curve fitting for the 15 min time point used weighting based on the standard deviation. For figure 5, curves are plotted as Δ anisotropy, where the basal fluorescence of the probe was subtracted from all points.

### Growth of yeast strains

Strains not requiring selection for auxotrophic markers (or requiring loss of a URA3 plasmid) were grown in standard YPDA (<u>Y</u>east extract/<u>P</u>eptone/<u>D</u>extrose (Glucose)/<u>A</u>denine) or YPD (without adenine) where indicated. Selection for URA3 strains was on SC-URA agar or broth (6.9g/L Yeast Nitrogen Base without amino acids (Formedium, CYN0405) + 0.96g/L Synthetic Complete Dropout - URA mixture (Formedium, DSCK1009) + 2% Glucose (Formedium, GLU03)). For CRAC, cells were grown in SMM-TRP (6.9g/L Yeast Nitrogen Base without amino acids + 740mg/L Complete Supplement Mixture - TRP (Formedium, DCS0149) + 2% Glucose). 5-FOA plates contained 6.7g/L Yeast Nitrogen Base without amino acids, 2% glucose, 20mg/L each of L-uracil, L-methionine and L-histidine, 50mg/L L-lysine and 100mg/L L-leucine + 1mg/ml 5-FOA (Formedium, 5-FOA01, dissolved at 100mg/ml in DMSO) and 2% Agar (Formedium, AGR05)).

### Construction of SSD1-tagged strains

All oligonucleotides and gBlocks were supplied by Integrated DNA Technologies (Supplementary File 2). Cloning strategies, designed using SnapGene (GSL Biotech LLC, 2365 Northside Dr., Suite 560 San Diego, CA 92108) are detailed in https://doi.org/10.5281/zenodo.4191152. To fuse the genomic copy of SSD1 with a C-terminal His-TEV-Protein A tag (HTP), a construct was designed (SSD1-HTP-URA3selplus) containing the last 100bp of the *S. cerevisiae* SSD1 ORF, an HTP-tag, a *Kluveromyces lactis* SSD1 3’UTR/Terminator and URA3 selection cassette and 100bp of the *S*.*cerevisiae* SSD1 3’UTR. The plasmid was synthesised by GeneArt (ThermoFisher), cut with SfiI, and integrated into the genome of BY4741 by homologous recombination after lithium acetate transformation (Gietz 2014). Colonies were grown on selective SC-URA plates, and two independent clones were verified by PCR.

Scarless integration of C-terminal HF-tags (HHHHHHHHAAAADYKDDDDK), or N-terminal FH-tags (DYKDDDDKAAAAHHHHHHHH), with and without deletion of the codons for the first 338 amino acids of SSD1, were made using a CRISPR/Cas9 yeast plasmid pML104 with URA selection (Laughery et al. 2015). Appropriate guide RNA (gRNA) sequences were identified using the CRISPR tools available at benchling.com, and imported into CRISPR tools courtesy of the Wyrick Lab (http://wyrickbioinfo2.smb.wsu.edu/crispr.html) to design oligonucleotides. These oligonucleotides were annealed and ligated into pML104 digested with SwaI and BclI, after growth in dam^−^ *E. coli*. Additional synonymous mutations within the guide RNA/PAM were included in the repair templates for the Ssd1-HF and FH-Ssd1 strains, to prevent further cleavage after repair. Repair templates (custom gBlocks; Integrated DNA Technologies) were amplified with Phusion Polymerase (New England Biolabs) for 12 cycles using specific gBlock-amplifying primers (Supplementary File 2). BY4741 yeast were transformed and selected as described above using 500ng of gRNA plasmid (URA3 selection) + /- 250-300ng of the relevant repair template. Clones were verified by PCR analysis and, sequencing. Once confirmed, tagged strains were grown overnight on non-selective medium (YPDA) and then plated on 5-FOA agar to select for loss of the gRNA plasmid.

### Yeast phenotyping assays

We investigated the sensitivity of our HTP and FH-tagged CRISPR strains compared to wild-type, ssd1Δ, and hsp104Δ, all in a BY4741 background, by growing them overnight to late log phase in 5ml YPD broth in culture tubes with vigorous shaking at 30°C.

For thermo-tolerance tests, 100ul of each overnight culture was transferred in duplicate to separate 200µl PCR tube strips. 1 strip was incubated for 30 mins at 37 °C and then cooled to 30 °C in a thermocycler. The second strip was incubated sequentially for 30 mins each at 37°C and then 50°C before cooling to 30 °C in a separate block in the same machine. Serial 5-fold dilutions in water were made into a 96-well plate and dilutions were replica plated on YPD plates, grown for 2 days at 30°C. For CFW sensitivity tests, 6x 10-fold serial dilutions were made in water of late log phase cultures of each strain. 5µl of each dilution was pipetted onto YPDA and YPDA+50µM CFW plates and grown at 30°C for 2 days.

### Cross-linking and analysis of cDNAs (CRAC) of Ssd1

A detailed protocol for the CRAC experiment is given at https://dx.doi.org/10.17504/protocols.io.5ppg5mn. In summary, two 2.86L cultures (in SMM-TRP medium in 5L flasks) for each of 2 biological replicates (independent clones) of the SSD1-HTP strains, plus one of BY4741 (untagged SSD1) as a control, were prepared from overnight pre-cultures at starting OD_600_ of approximately 0.05, and shaken at 30°C until they reached an OD_600_ of 0.45. Each replicate of the SSD1-HTP strain was filtered rapidly through 0.45µM nitrocellulose membrane filters (Millipore, HAWP09000) to collect the cells. One set of filtered cells from each biological replicate was transferred on the membranes to 5L flasks containing 2.86L of SMM-TRP medium pre-warmed to 42°C and shaken at 42°C for 16 mins before immediate transfer of the cultures to the Megatron (UVO3; (Bohnsack, Tollervey, and Granneman 2012)) for UVC (254nm) cross-linking for 100s. Cells were recovered again by filtration, washed in water and transferred to 30ml of PBS in a 50ml Falcon tube, shaking to release the cells, removing the membranes, pelleting the cells and draining the tubes before storing at −80°C. The remaining cultures were taken straight from 30°C for cross-linking and downstream treatment as above. Extracts of the cross-linked pellets were processed into sequencing libraries as previously described (Bohnsack, Tollervey, and Granneman 2012) using 1µl of a 1:100 dilution of 10U/µl RNace-IT (Agilent Technologies, 400720) per sample, 22 cycles of PCR after Reverse Transcription and size selection of products of around 120-180bp (average 150bp), for full details see protocols.io. Library concentrations were measured using the Qubit ds HS Assay Kit (Q32851) and pooled at 1nM final concentration for SE (<u>S</u>ingle <u>E</u>nd) read sequencing with an Illumina MiniSeq High Output Reagent Kit (75-cycles, FC-420-1001) on an Illumina MiniSeq System Instrument.

### CRAC Data Analysis

The complete pipeline and intermediate data for CRAC data analysis, including figure construction, is available at https://doi.org/10.5281/zenodo.4191152. In brief, we adapted the single end reads pipeline developed by the Granneman lab (van Nues et al. 2017), relying on multiple tools from the PyCRAC software suite (Webb et al. 2014). Initially, the 3’adapters were removed from the FASTQ files using flexbar and then pyBarcodeFilter.py was used to demultiplex the fastq files based on their barcodes. pyFastDuplicateRemover.py was used to collapse PCR duplicates based on identity of both the insert sequence and the random nucleotides in the barcodes. Collapsed FASTA files were then aligned to the yeast genome using Novoalign 2.0 (Novocraft technologies). Reads were counted using multiBamCov from bedtools (Quinlan 2014), to transcript maps from Saccharomyces Genome Database (Ng et al. 2020), using the “abundant transcript” data derived from (Pelechano, Wei, and Steinmetz 2013), adding default-length 25nt 5’UTRs and 125nt 3’UTRs for verified ORFs missing from that annotation. Bedgraph files were generated using genomeCoverageBed from bedtools (Quinlan 2014). Pileup files, including deletions and mutations, were made using pyPileup.py running on selected Ssd1-associated transcripts. Count output gtf files were made using pyReadCounters.py, then pyCalculateFDRs.py was used to detect enriched peaks with a False Discovery Rate ≤ 0.05. We filtered to the top 100 peaks by height and searched for enriched motifs using MEME (Bailey et al. 2015). RNA-seq data from GEO: GSE148166 was similarly aligned using Novoalign 2.0 and assigned to the same transcripts with multiBamCov from bedtools (Quinlan 2014). Data were further analysed and visualised the in R (Team 2020), using ggplot2 (Wickham 2016), tidyverse packages (Wickham and 2019) and R markdown (Xie 2018).

### Evolutionary analysis

For alignment of Ssd1 binding site on SUN4 5’UTR in *Saccharomyces*, orthologues of SUN4 in *S. cerevisiae, S. paradoxus, S. mikatae, S. kudriavzevii, S. arboricola, S. uvarum*, and *S. eubayanus* were selected and sequences 700 nt upstream of the start codon were retrieved using orthology mapping and annotations provided by (Shen et al. 2018). These were aligned using MAFFT v7.429, option *genafpair* (Katoh and Standley 2013), and the sequence logo was computed with ggseqlogo (Wagih 2017). Ascomycete homologues were chosen from PANTHER family PTHR31316:SF0 (Mi et al. 2019), and their protein and transcript annotations were obtained from FungiDB (Basenko et al. 2018). The annotation of the 5’UTR of *C. albicans* SUN41 was adjusted to account for its 5’UTR intron (Mitrovich et al. 2007). Motif occurrences were counted by eye and where overlapping sequences such as CNYUCNYUCNYU were observed these were counted as 2 occurrences of CNYUCNYU. For protein phylogeny, we aligned the sequences using MAFFT v7.429, option genafpair (Katoh and Standley 2013), computed the tree with fasttree 2.1.10 (Price, Dehal, and Arkin 2010), and plotted the figure using ggtree (Yu et al. 2018).

### Data sharing

Coordinates for the Ssd1 structure were deposited in the PDB, PDBid 7AM1. CRAC datasets have been deposited on GEO, accession numbers GSE159835.

## Acknowledgements

We thank Sutapa Chakrabarti, Elena Conti and Elizabeth Blackburn for reading and commenting on the manuscript. This work was supported by the Wellcome through a Senior Research Fellowship to AGC (200898), a Sir Henry Dale Fellowship with The Royal Society to EW (208779), a Principal Research Fellowship to DT (109916), a Centre Core grant to the Wellcome Centre for Cell Biology (203149), and ISSF funding. The authors would like to thank Diamond Light Source for beamtime (proposal mx18515), and the staff of beamline I04-1 and A. A. Jeyaprakash for assistance with data collection. We thank the staff of the Edinburgh Protein Production facility for access to protein purification and characterisation equipment. We thank Sander Granneman with help on pyCRAC software and CRAC data analysis. We thank Shaun Webb and Hywel Dunn-Davies at the Wellcome Centre for Cell Biology bioinformatics core facility.

## Author contributions

AGC and EWJW conceived and directed the research. UJ purified and crystallized the protein. AGC did the crystallographic data analysis. RAB created the CRISPR tagged strains and did phenotyping analysis. RAB and SB carried out CRAC sample and library preparation. EWJW and RAB analysed CRAC and RNA-seq data. AK carried out EMSA and FA analysis. AGC, EWJW and DT wrote and edited the manuscript, with assistance from all authors.

