## Supplementary figures for "Yeast Ssd1 is a non-enzymatic member of the RNase II family with an alternative RNA recognition interface"

#### Supplementary Figure 1

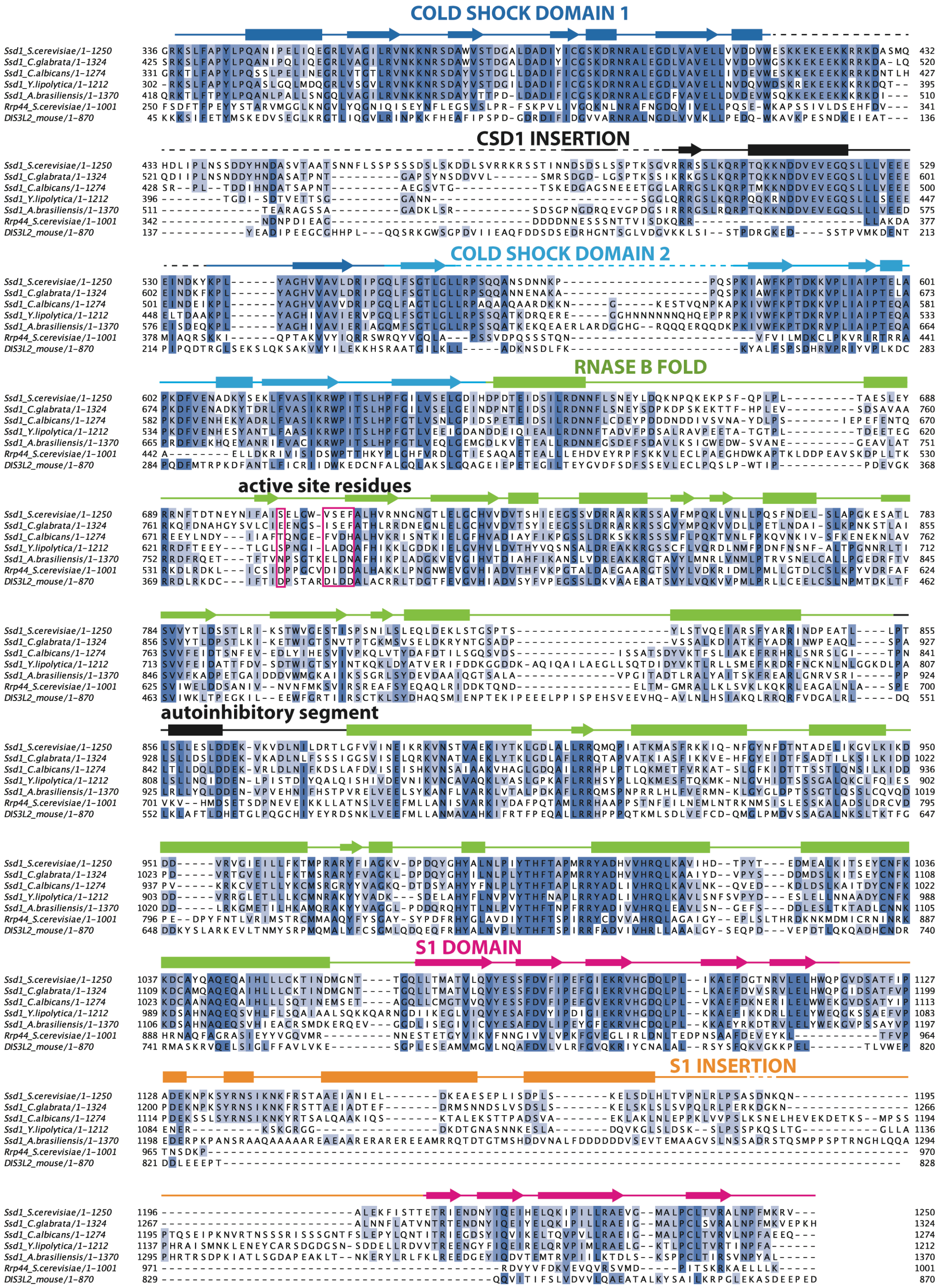


**Figure S1.** A multiple sequence alignment that includes several Ssd1 homologues from Fungi as well as yeast Rrp44 and mouse Dis3L2. The alignment is coloured based on percent identity. Secondary structure elements for alpha helices (rectangles) and beta strands (arrows) are shown above the sequence, with unassigned segments represented by dotted lines. Pink boxes indicate residues equivalent to active site signature in active RNB family enzymes.

### Supplementary figure 2

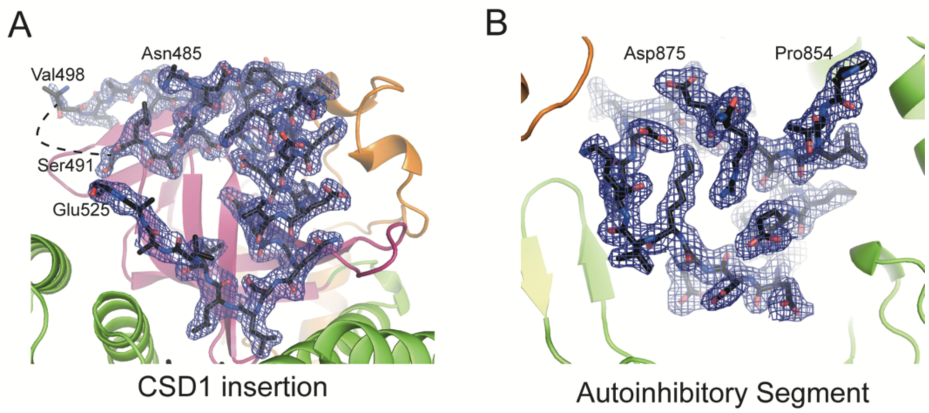


**Figure S2.** (**A**) Electron density around the CSD1 insertion. The view is the same as the left view in Figure 1B but with the two CSD domains removed for clarity. (**B**) Electron density around the autoinhibitory segment. The view is the same as the right-hand view in Figure 1B but rotated through 90° around the x-axis, for clarity.

### Supplementary figure 3

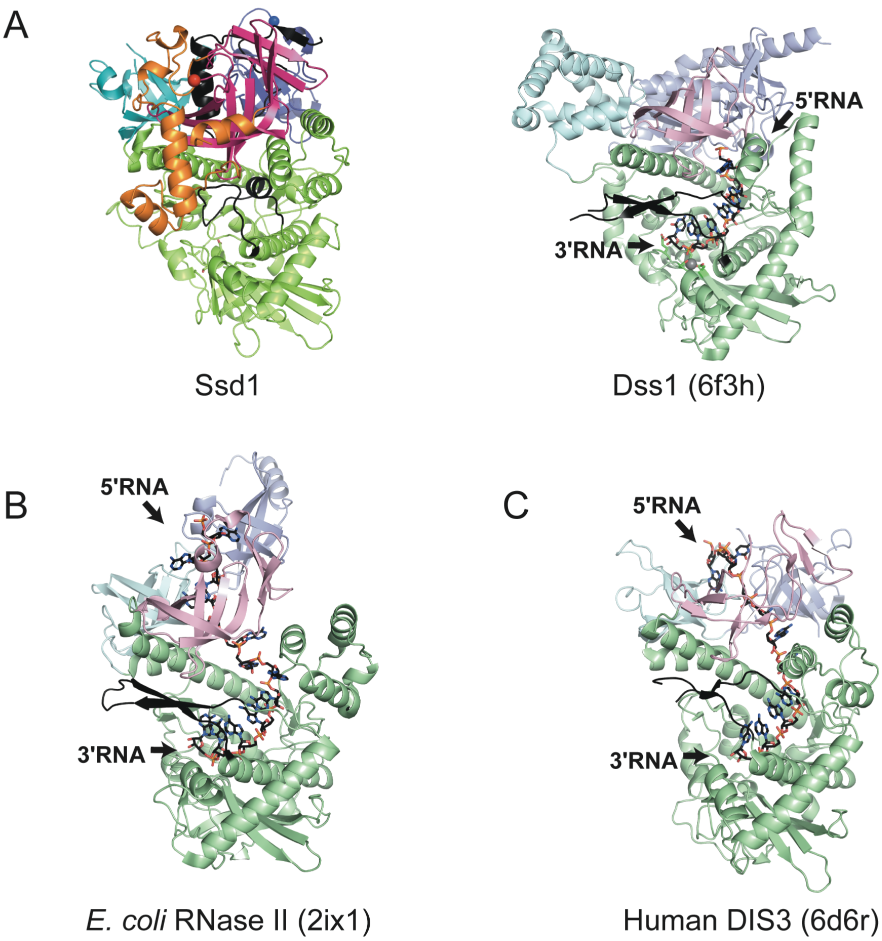


**Figure S3.** (**A**) Ssd1 is compared by superposition with RNA-bound Dss1 (PDBID 6f3h). (**B**) *E. coli* RNase II (PDBID 2ix1) bound to RNA is shown in the same orientation of Ssd1 as in (**A**). (**C**) Human DIS3 (PDBID 6d6r) from the exosome structure containing Mtr4 and RNA is shown in the same orientation as (**A**). The exosome subunits and part of the RNA are removed for clarity.

Supplementary figure 4

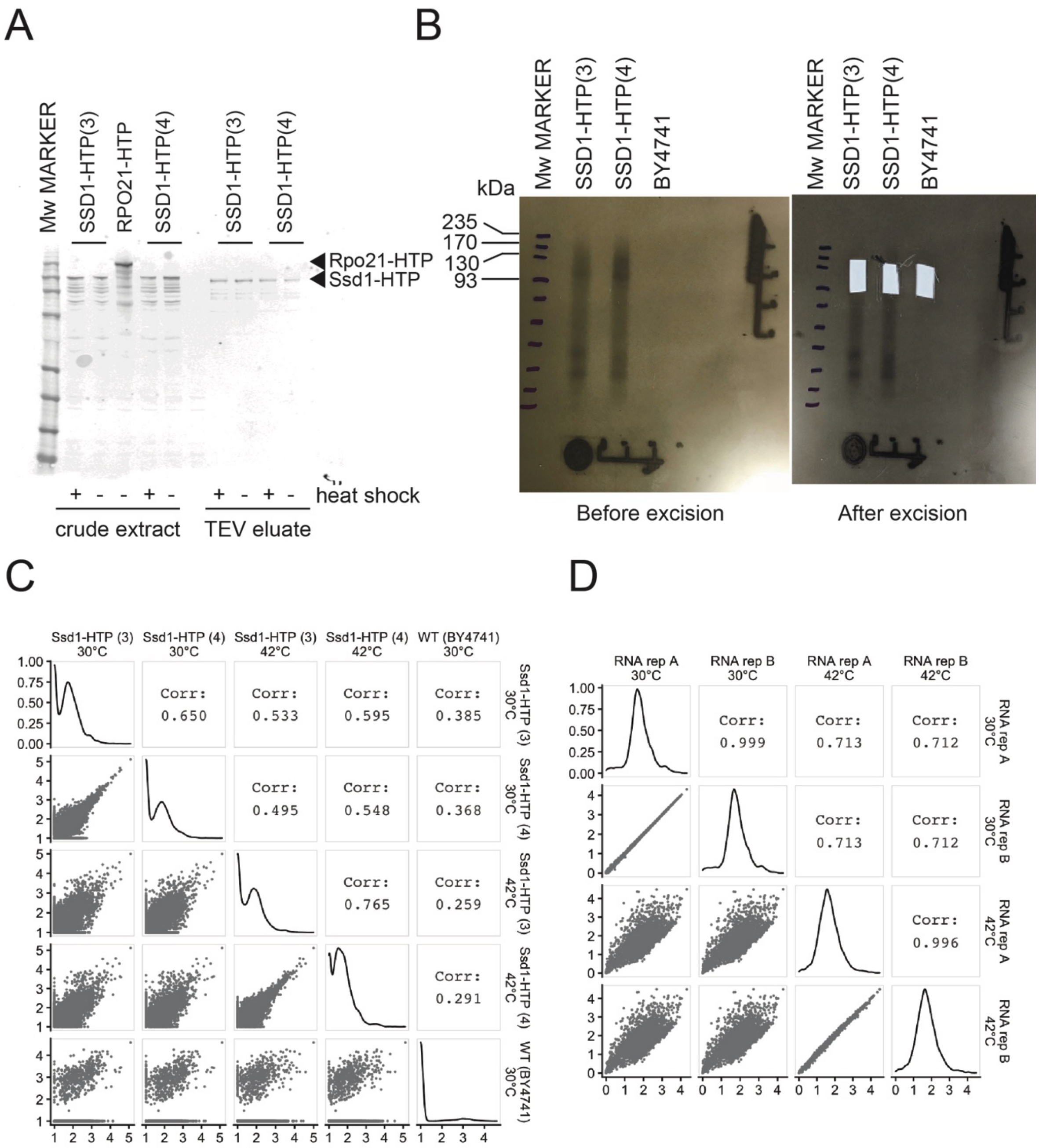


**Figure S4.** (**A**) Western blot of the crude extracts and TEV eluted CRAC samples was probed with rabbit anti-TAP antibody (ThermoFisher CAB1001; 1/5000 dilution) followed by donkey anti-rabbit Dylight680 antibody (ThermoFisher SA5-10042; 1/10000 dilution) and imaged on a LICOR Odyssey CLx Infrared Imaging System. (**B**) Autoradiograph of $^{32}$P - labelled SSD1-RNA complexes before and after extraction of the smear above the Ssd1 protein used to prepare the libraries. (**C**) Pairwise comparison of read densities on verified coding transcripts from CRAC datasets, in units of log10 of transcripts per million (TPM). (**D**) Pairwise comparison of read densities on verified coding transcripts from RNA-seq datasets, also in log10(TPM).

#### Supplementary figure 5

A

| Meme | LOGO | E value | No. of nucleotides | No. of sites |
|---|---|---|---|---|
| 1 | 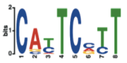 | 2.4e-037 | 8 | 59 |
| 2 | 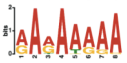 | 9.8e-010 | 8 | 59 |
| 3 | 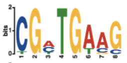 | 2.9e-004 | 8 | 15 |
| 4 | 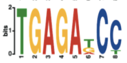 | 1.5e-001 | 8 | 9 |
| 5 | 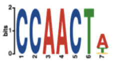 | 7.5e-001 | 7 | 16 |

B

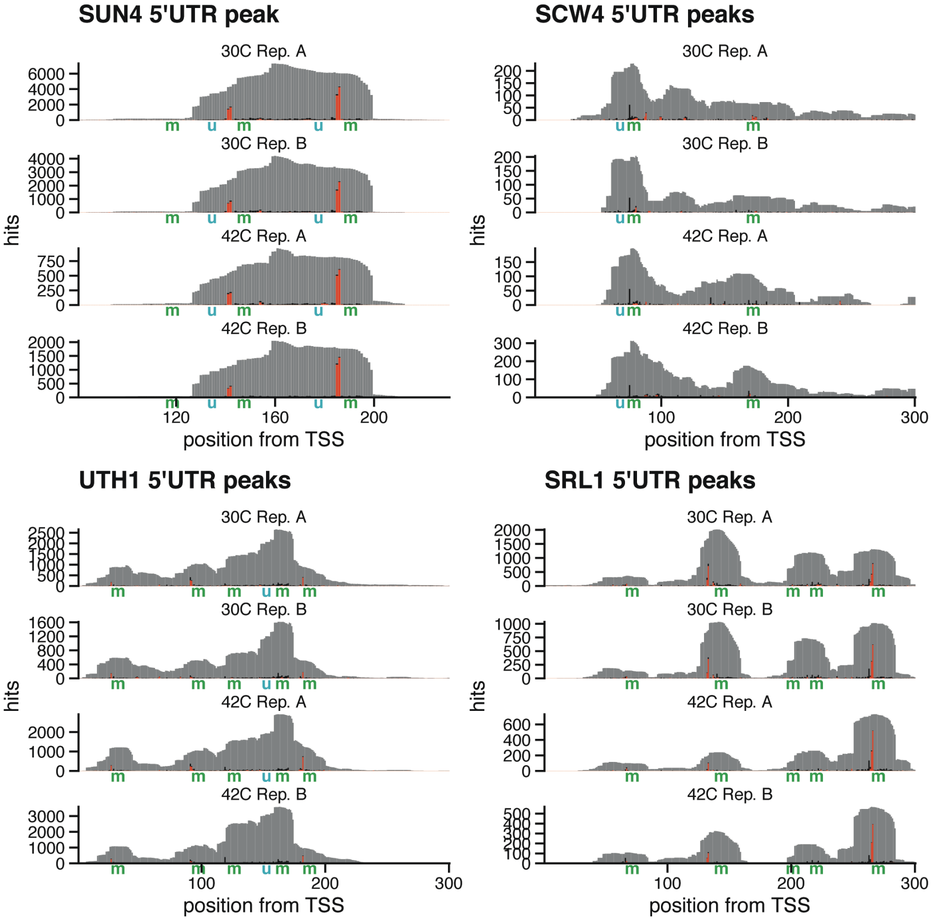


**Figure S5**. (**A**) Logos of sequence motifs enriched in transcripts around CRAC crosslink sites. The top five logos from MEME analysis are shown. (**B**) Pileup plots of CRAC hits which match the genome (grey) or have mutations (black) or insertions (red), showing all 4 replicates on 5' ends of four selected transcripts, SUN4, SCW4, UTH1, and SRL1. CNYUCNYU positions are marked with "m" for motif, and CCAACU positions with "u" for upstream.

### Supplementary figure 6

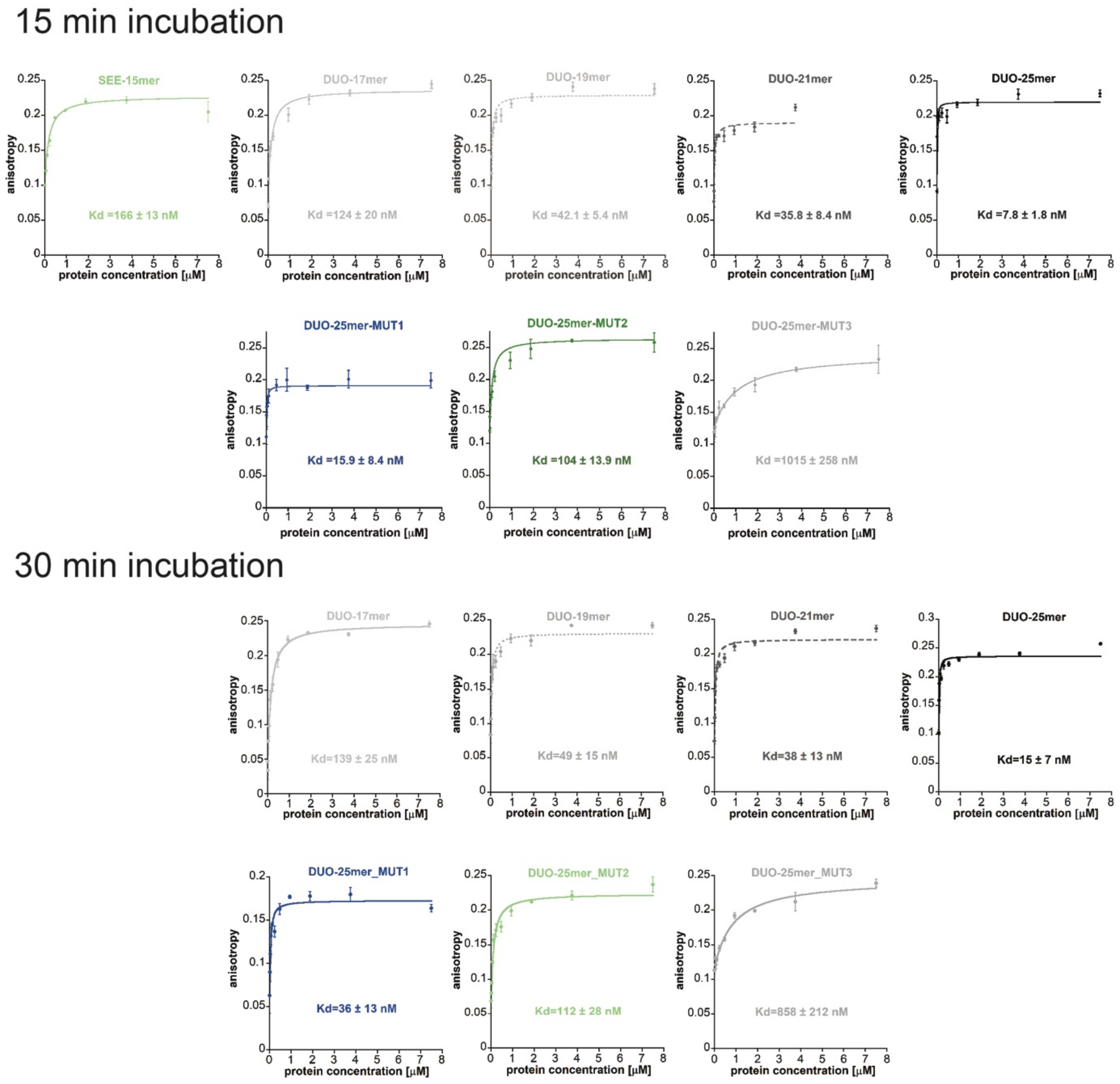


**Figure S6.** Fluorescence anisotropy data used in Fig. 5 shown as individual experiments with fitted curves. Data from an additional set of experiments carried out with longer incubation times are included for comparison. Graph fitting for measurements at 15 mins were weighted by standard deviation.
